# A Bayesian Non-parametric Mixed-Effects Model of Microbial Phenotypes

**DOI:** 10.1101/793174

**Authors:** Peter D. Tonner, Cynthia L. Darnell, Francesca M.L. Bushell, Peter A. Lund, Amy K. Schmid, Scott C. Schmidler

## Abstract

Substantive changes in gene expression, metabolism, and the proteome are manifested in overall changes in microbial population growth. Quantifying how microbes grow is therefore fundamental to areas such as genetics, bioengineering, and food safety. Traditional parametric growth curve models capture the population growth behavior through a set of summarizing parameters. However, estimation of these parameters from data is confounded by random effects such as experimental variability, batch effects or differences in experimental material. A systematic statistical method to identify and correct for such confounding effects in population growth data is not currently available. Further, our previous work has demonstrated that parametric models are insufficient to explain and predict microbial response under non-standard growth conditions. Here we develop a hierarchical Bayesian non-parametric model of population growth that identifies the latent growth behavior and response to perturbation, while simultaneously correcting for random effects in the data. This model enables more accurate estimates of the biological effect of interest, while better accounting for the uncertainty due to technical variation. Additionally, modeling hierarchical variation provides estimates of the relative impact of various confounding effects on measured population growth.

## 1 Introduction

Population growth phenotypes inform studies in microbiology, including gene functional discovery, bioengineering process development, and food safety testing^1–3^. For example, recent advances in microbial functional genomics and phenotyping, or “phenomics”, have enabled transformative insights into gene functions, proving critical for mapping the genotype to phenotype relationship^4^. Methods such as genome-wide CRISPRi^5^ and targeted genome-scale deletion libraries^6,7^ frequently rely upon accurate quantitation of microbial population growth as an assay to identify novel mutants with significant growth phenotypes. Population growth is an aggregate measure of all cellular processes and captures how microbial cells adapt and survive in their environmental niche^8^. Because microbial population culturing is a necessary precursor to many experimental procedures in microbiology^9^, reproducible results require accurate quantification of the variability in culture state measured through growth^9,10^.

Typical analyses of microbial population growth involve estimating parametric models under the assumptions of standard growth conditions comprised of three successive growth phases: (1) lag phase, in which the population adapts to a new environment, typically fresh growth medium at culture inoculation; (2) log phase, when the population grows exponentially at a rate dependent on nutrients in the environment; and (3) stationary phase, where measurable population growth terminates thereby reaching the culture carrying capacity^11^. Recent studies have shown that the estimates of parameters in these models are highly uncertain^12–14^. This uncertainty arises both from factors of biological interest, such as differences in genetic background and environment, as well as uncontrolled technical noise from experimental manipulation of microbial cultures. While such sources of variability can be modeled using fixed and random effects^15–19^, parametric population growth models have additional limitations. Most notably, when population growth deviates from the standard sigmoidal shape assumed in parametric models, secondary models must be developed on a case by case basis for each new experimental perturbation^20,21^. Additionally, we have shown in previous work that in cases such as extreme stress or strongly deleterious mutations, no parametric growth model accurately represents the growth curve, regardless of secondary model^19,22,23^.

Factors affecting microbial growth measurements include both fixed and random effects^24^. Fixed effects are assumed to be drawn from a finite set of perturbations of interest, for example the effect of different concentrations of a chemical on growth that are entirely represented in the dataset. Random effects, conversely, can be viewed as a random sample from a larger population of interest. For example, repeating the same design over many experiments corresponds to sampling the random experimental effect from the theoretical population of all possible experiments that could be conducted with this design^3,25^. Random effects arising from repeated experimental design are typically referred to as *batch effects*^26,27^. Batch effects are often a significant component of measurement noise in high-throughput genomics experiments^28^. However, random effects are not always due to experimental noise, and may represent quantities of direct scientific interest; for example, assaying a set of genetic backgrounds may be viewed as sampling from the population of all possible genetic variants^29–33^. Models which include both fixed and random effects are referred to as mixed effects models.

In this study we present *phenom*, a general model for analysis of phenomic growth curve experiments based on a Bayesian non-parametric functional mixed effects model of microbial growth. We demonstrate the utility of *phenom* model to analyze population growth measurements of two microorganisms: the hypersaline adapted archaeon, *Halobacterium salinarum*; and the opportunistic bacterial pathogen, *Pseudomonas aeruginosa. H. salinarum* is a model organism for transcriptional regulation of stress response in the third domain of life, the Archaea^34–36^. *H. salinarum* is particularly well adapted to resisting oxidative stress (OS), which arises from the buildup of reactive oxygen species and causes damage to many critical cellular components, including DNA, protein, and lipids^37–43^. Population growth measurements of *H. salinarum* under OS have been used previously to quantify these harmful effects on physiology, as well as identify regulatory factors important for OS survival^22,40–42^. The presence of batch effects in *H. salinarum* OS response was reported (and corrected for) previously^19^, but did not model individual batch effects for each term in the model. This motivated the explicit deconstruction of batch effects between different factors (e.g. strain and stress), which we have implemented and reported here in *phenom*.

*Pseudomonas aeruginosa* is an opportunistic microbial pathogen and a growing problem in hospital-borne infections. Rising antimicrobial resistance of these organisms has necessitated the development of alternative treatment strategies. For example, topical treatment of infected burn wounds with acetic or organic acids (OAs) has been successful^44^. OA impact on growth depends on external pH levels — in acidic intracellular environments the OA does not dissociate, freely traverses the cellular membrane as an uncharged particle, and dissociates in the neutral cytoplasm inducing acid stress^45^. Here we apply *phenom* to the *P. aeruginosa* dataset, which is foundational for a larger study of *P. aeruginosa* strains responding to pH and OA perturbation as a potential novel treatment of pathogenic bacterial infections^23^.

Stress occurs constantly in the environment: as conditions change, mild to severe cellular damage occurs, and cells must regulate their molecular components to survive^46–49^. Population growth measurements are particularly vital to the study of stress response by providing a quantitative measure of growth differences against a non-stressed control^1^. Our model recovers fixed effects due to high and low levels of oxidative stress in *H. salinarum* as well as interactions between organic acid concentration and pH in *P. aeruginosa*, while correcting for random effects from multiple sources, thus enabling more accurate estimates of the significance of the stress treatment effect. Notably, in cases where random effect and fixed effect sizes are comparable, we demonstrate that mixed modeling is critical for accurate quantification of model uncertainty. If random effects are not included in the model, the significance of the effect of stress treatments on population growth can be erroneously overestimated. We discuss the implications of these findings for multiple areas of microbiology research.

## 2 Results

### 2.1 Hierarchical batch effects typical in phenomics datasets render parametric models ineffective

In the dataset used here, population growth for each of *P. aeruginosa* and *H. salinarum* cultures was monitored under standard (non-stressed) conditions vs. stress conditions (see Materials and Methods and references [22, 23] for precise definition of “standard conditions” for each organism). Specifically, cultures were grown in liquid medium in a high throughput growth plate reader that measured population density at 30 minute intervals over the course of 24 hours (*P. aeruginosa*) or 48 hours (*H. salinarum*); the resulting data are shown in Fig. 1. Experimental designs for each organism included biological replicates (growth curves from different colonies on a plate), technical replicates (multiple growth curves from the same colony), varying conditions (stress vs standard), and are further divided into batches (different runs of the high throughput growth plate reader). *H. salinarum* was grown under high (0.333 mM paraquat (PQ)) and low (0.083 mM PQ) levels of oxidative stress (OS); the data are combined from published^19,22,41^ and unpublished studies (Fig 1A). The OS responses of *H. salinarum* were compared to a control of standard growth in rich medium, representing optimal conditions for the population. The experimental design was replicated in biological quadruplicate and technical triplicate, across nine batches (Fig. 1A, individual curves and axes). *P. aeruginosa* was grown in the presence of increasing concentrations of three different organic acid (OA) chemicals (0 — 20mM; benzoate, citric acid, and malic acid), each combined with a gradient of pH (5.0 — 7.0)^23^. Each *P. aeruginosa* growth condition was repeated across 3 biological replicates and two batches (Fig 1B). The different *P. aeruginosa* and *H. salinarum* experimental designs with varying numbers of replicates at each level provides a rich testbed for exploring the impact of modeling random effects with *phenom* (Figs. 1B, S1, S2).

**Figure 1:**
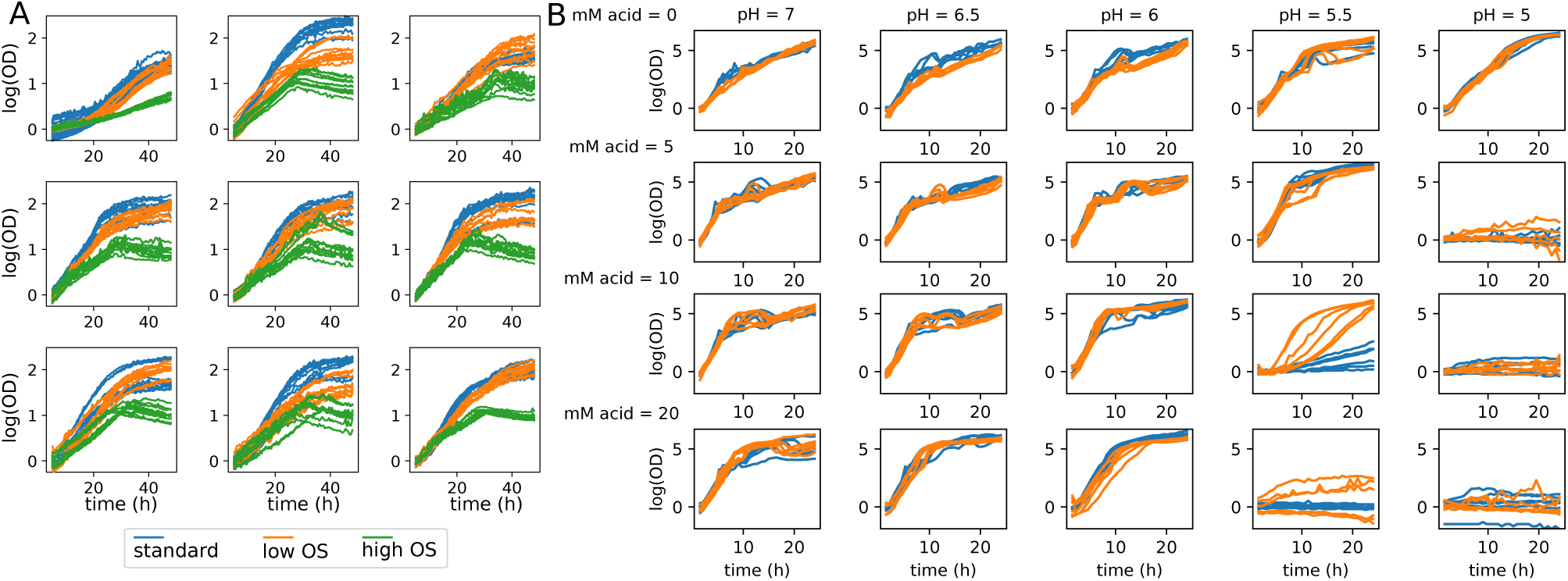
Batch variation in high throughput phenomics studies. (A) Population growth measurements of *H. salinarum* under standard conditions (blue), and low (orange) and high (green) levels of OS. Individual measurement curves are replicates and each graph panel is a different batch. (B) Growth of *P. aeruginosa* strain PA01 under gradient of pH (5 — 7) and citric acid (0 — 20 mM). Colors represent different batches.

Figures 1 and 2 demonstrate the two key issues described above and addressed in this paper. First, batch effects are present in both *H. salinarum* and the *P. aeruginosa* datasets. For *H. salinarum*, clear differences in growth under both standard and stress conditions are observed in the raw data across experimental batches (i.e. separate runs of the growth plate reader instrument; Fig. 1). Some batches show a different phenotype, with either a complete cessation of growth or an intermediate effect with decreased growth relative to standard conditions. For example, in some batches, populations stressed with low OS grow at the same rate and reach the same carrying capacity as populations grown under standard conditions. For *P. aeruginosa*, a clear difference between batches grown under 10 mM citric acid at pH=5.5 is observed [Fig. 1B (graph in fourth column, third row) and Fig. 2D]. Like with citric acid, batch effects were also found in some of the other conditions considered (e.g. growth under malic acid, Figs. S1, S2).

**Figure 2:**
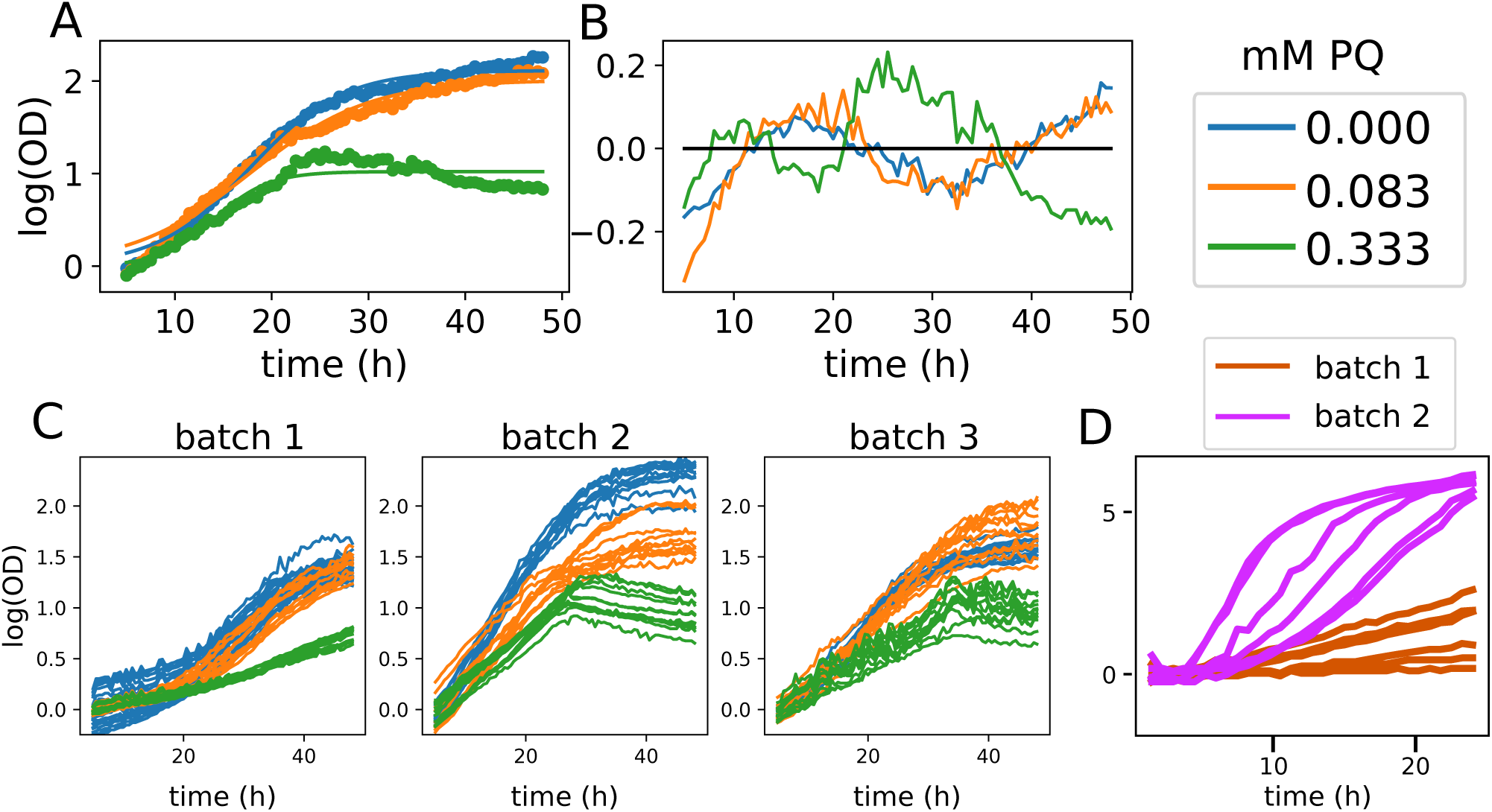
Batch effects are prevalent in microbial phenomic datasets. (A) Parametric fits to *H. salinarum* growth curves. (B) Residuals of parametric growth curve fit. (C) Growth of *H. salinarum* under standard conditions (blue), low (orange) and high (green) OS across three batches. (D) Measurement of *P. aeruginosa* growth under 10mM citric acid at 5.5 pH. Measurements for each condition vary significantly with batch.

Second, standard parametric growth curve models fail to describe experimental measurements adequately (Fig 2A, B), as we have shown previously with both datasets^19,22,23^. In Fig. 2, we examined the impact of batch and replicate effects on our data by considering how they change parameters estimated from a mixed effects parametric model of population growth^32^. We focused on calculating *µ*_max_, the maximum instantaneous growth rate attained by the population, as this is a commonly used parameter for comparisons between conditions^19,50^. Variation in *µ*_max_ estimates were observed both on the replicate and batch level, as shown by the kernel density estimates (KDE) of *µ*_max_ for each stress level (Fig. S3). The variance in *µ*_max_ is remarkably high: the 95% confidence interval for *µ*_max_ under standard growth is 0.050—0.141, a nearly 3-fold change between the lower and upper interval limits. Thus, while the t-test conducted on *µ*_max_ estimates between standard conditions and each stress level is statistically significant (Fig S3), it is difficult to conclude: (a) what the true magnitude of the stress effects may be; and (b) to what degree the variation due to replicate and batch should inform biological conclusions. The error of the logistic growth model under each PQ condition was also examined. Error increased under high OS (Fig S4). High OS induces a growth phenotype that deviates heavily from the sigmoidal growth curve assumed in the logistic model as well as in other commonly used growth models. This leads to a poor fit under the high OS condition as has been shown previously (Fig S4 ^19^). The residuals under standard, low, and high OS conditions also appear to be dependent. Our previous work also demonstrated poor fits to the *P. aeruginosa* data using parametric models^23^. Taken together, the initial assessment of these two datasets indicates that: (a) technical variation due to batch and replicate in growth curve data can be high; and (b) commonly used standard parametric models are not able to adequately capture or correct for these sources of variability. These sources of error need to be corrected in order to model true growth behavior and inform biological conclusions from the data.

### 2.2 A hierarchical Bayesian model of functional random effects in microbial growth

We previously established the ability of non-parametric Bayesian methods to improve the modeling of growth phenotypes^19,22,23^. Here, we describe *phenom*, a fully hierarchical Bayesian non-parametric functional mixed effects model for population growth data. We highlight the utility of *phenom* to correct for confounding, random effects in growth phenotypes.

In order to model both biological and technical variation in microbial growth (Fig 3), we first assume that a set of population growth measurements are driven by an (unobserved) population curve *µ*(*t*) (Fig 3A, blue curve) of unknown shape. For example, *µ*(*t*) might represent the average growth behavior of an organism under standard conditions. This mean growth behavior may be altered by a treatment effect, represented by an additional unknown curve *δ* (*t*) (Fig 3A, orange curve). For example *δ* (*t*) may represent the effects on growth induced by low or high levels of OS (Fig 2A). The average growth behavior of a population under stress conditions would then be described by the curve *f* (*t*) = *µ*(*t*) + *δ* (*t*).

**Figure 3:**
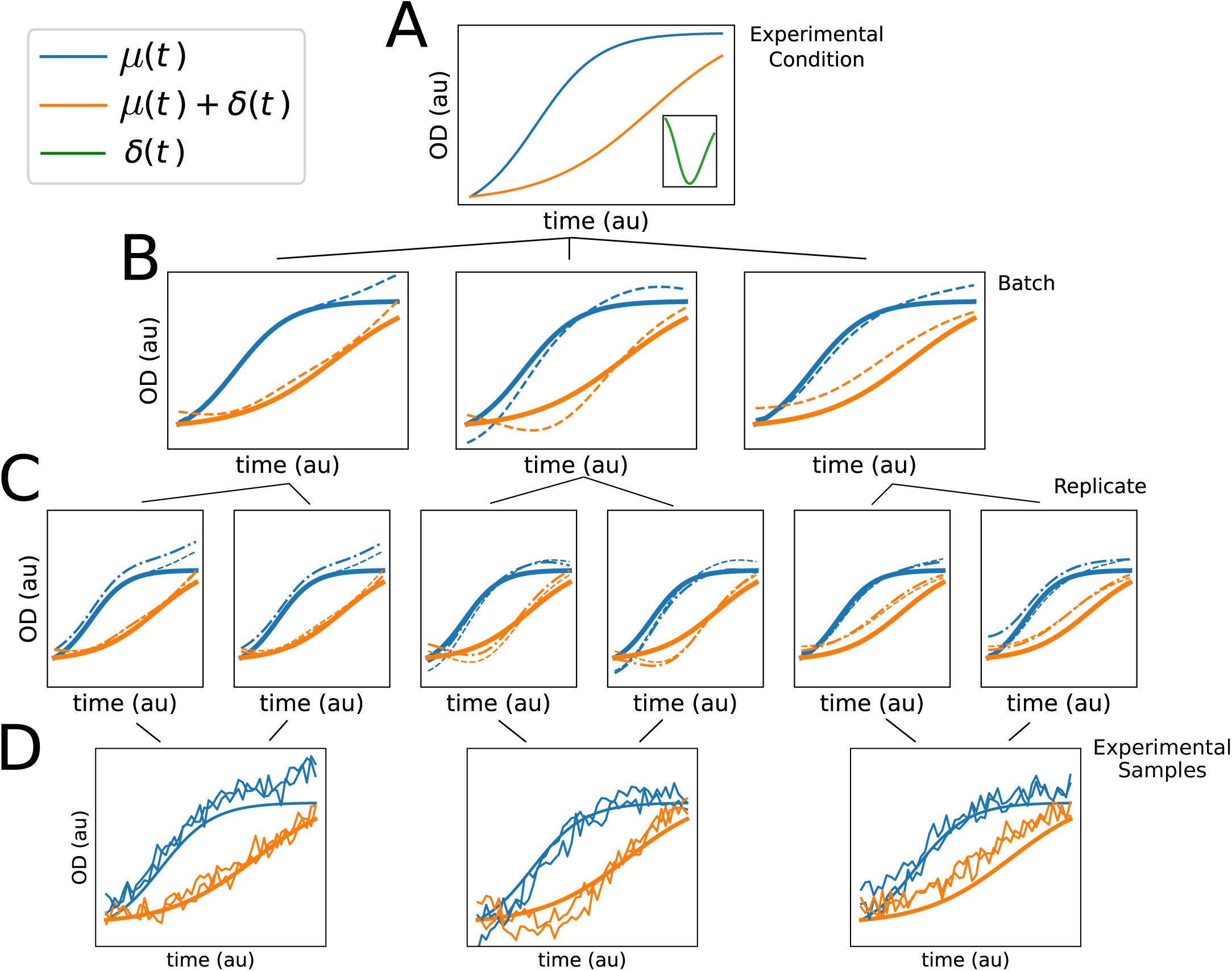
Hierarchical model of functional data. Representative diagram of hierarchical variation present in microbial growth data. Each tier of graphs represents a different variation source, and lines indicate relationship between them: experimental condition is the true growth behavior of interest, with the condition repeated across batches, and replicates repeated within each batch. (A) Functional phenotypes *µ*(*t*) (blue), *µ*(*t*) + *δ* (*t*) (orange), and *δ* (*t*) (green curve in inset). (B) Batch effects on *µ*(*t*) and *µ*(*t*) + *δ* (*t*). Each plot is a different batch, solid lines are the true functions as in (A), and the dashed lines are the observed batch effect of *µ*(*t*) and *µ*(*t*) + *δ* (*x*) for the corresponding batch. (C) Replicate effect within batches. Each axis is a different replicate, solid and dashed lines as in (B), dotted-dashed line is the observed replicate function. (D) Observations from the model described in (A-C). Each curve is sampled with a mean drawn from the global mean, with added batch and replicate effects (dotted-dashed lines in C) and *iid* observation noise. Each axis is a different batch. The smooth solid lines are the true functions *µ*(*t*) and *µ*(*t*) + *δ* (*t*) in (A).

When considering a combinatorial experimental design, such as that described for *P. aeruginosa* growth (Fig. 1B), we model independent effects of different treatments as well as their interaction via the form:

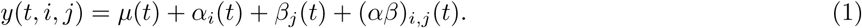

Here, *y*(*t, i, j*) denotes the observed population size at time *t* with treatments *i* and *j* of two independent stress conditions. Additionally, *α*_*i*_(*t*) and *β*_*j*_(*t*) are the independent effects of each stress condition, and (*αβ*)_*i,j*_(*t*) is their interaction. This model corresponds to a functional analysis of variance^51^, which we have previously used to estimate independent and interaction effects of microbial genetics and stress^22^. Here, we consider the interaction of the two stress conditions as well as random functional effects in the model.

Variability around these fixed effect growth models is described by additional, random curves associated with two major sources of variation: *batch* and *replicate* (Fig 3B,C). Batches correspond to a single high-throughput growth experiment and replicates are the individual curve observations within a batch. Using *phenom* throughout this study, we only compare replicates that are contained within the same batch. This is due to the nested structure between batch and replicates (Fig 3). Noise due to both replicate and batch do not appear to be independent identically distributed (*iid*), as observed in the correlated residuals around the mean for each experimental variate (Fig. S5A and B). Each observed growth curve is therefore described by a combination of the fixed effects and the corresponding batch and replicate effects (Fig 3D). Both replicate and batch variation are modeled as random effects because the variation due to both sources cannot be replicated, i.e. a specific batch effect cannot be purposefully re-introduced in subsequent experiments. Instead, these variates are assumed to be sampled from a latent super-population^52^. Combining the fixed and random effects, we arrive at a mixed-effects model of microbial phenotypes.

We adopted a hierarchical Bayesian framework to model these mixed effects. In this framework, batch effects are described by a shared generative distribution, allowing them to take on distinct values while still pooling across replicates for accurately estimating the generating distribution^53^. We use Gaussian process (GP) distributions for all groups in the model. GPs are flexible, non-parametric distributions suitable for smooth functions^54^. To assess the impact of incorporating random effects on estimation of the treatment effect of interest, we analyze three models of increasing complexity: M_null_ excludes all hierarchical random effects, M_batch_ incorporates batch variation only, and M_full_ incorporates both batch and replicate variation. These models, collectively called *phenom*, were implemented using the probabilistic programming language Stan^55^, which efficiently traverses the posterior through Hamiltonian Monte Carlo (see Materials and Methods).

In order to demonstrate the impact of batch effects on the conclusions drawn from the analysis of microbial growth data, we estimated the latent functions driving both *H. salinarum* and *P. aeruginosa* growth using the M_null_ model of *phenom*, with each batch analyzed separately (Fig 4). This corresponds to the analysis that would be conducted after generating any single set of experiments from a batch, without considering or controlling for batch effects, and therefore provides a test of the impact of ignoring batch effects.

**Figure 4:**
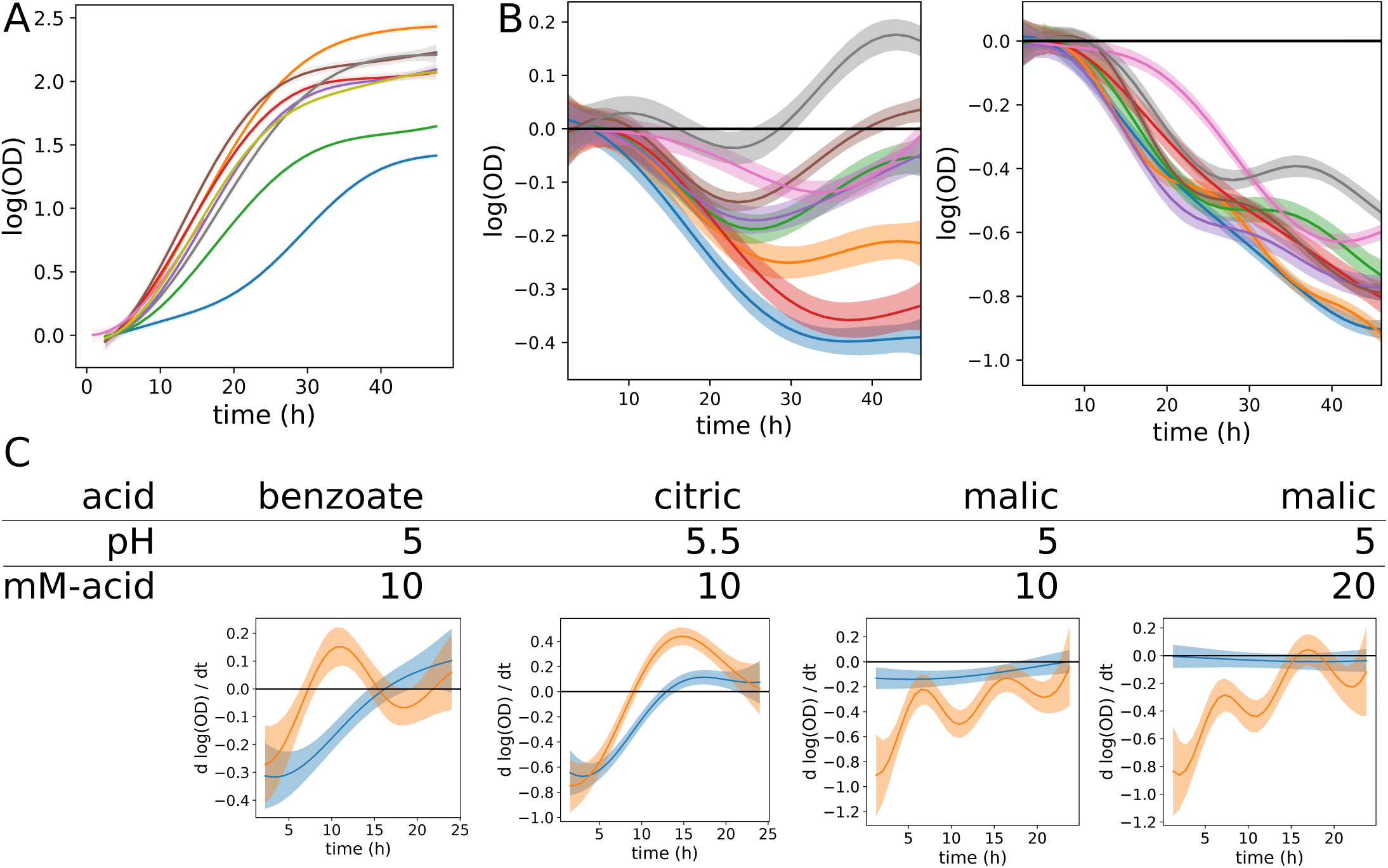
M_null_ model estimates are confounded by batch effects. Posterior intervals of functions are shown for different analyses where *phenom* M_null_ was fit using data from each batch separately. In all plots, solid line represents posterior mean, shaded region indicates 95% credible region, and each color corresponds to a different posterior conditioned on data from a single batch. (A) Posterior intervals of *µ*(*x*), the standard growth phenotype of *H. salinarum*. (B) Posterior interval of *δ* (*x*) under low (left) and high (right) OS response of *H. salinarum*. (C) Posterior interval of interaction function (*αβ*)_*p,m*_(*t*) for *P. aeruginosa* growth in indicated pH and acid concentration.

For *H. salinarum*, growth data under standard conditions was used to estimate a single mean function, *µ*(*t*) (Fig. 4A). Fixed effects for growth under low and high OS was added as *δ*(*t*) (Fig 4B). For the *P. aeruginosa* dataset, batch effects on the interaction between pH and organic acid concentration was represented by a function (*αβ*)_*p,m*_(*t*), again estimated non-parametrically (Fig. 4C). However, rather than reporting (*αβ*)_*p,m*_(*t*) directly, we report its time derivative, which has the interpretation of instantaneous growth rate rather than absolute amount of growth^56^.

Fitting the M_null_ model to each separate batch reveals that the posterior distributions obtained for each function of interest (*µ*(*t*), *δ* (*t*), and (*αβ*)_*p,m*_(*t*)) are highly variable across batches (Fig. 4). This is observed in both the *H. salinarum* and *P. aeruginosa* datasets, where the experimental conditions, and therefore the underlying true functions, remain constant across batches in each case. Such variability can impact conclusions. For example, in the low OS condition in the *H. salinarum* dataset, both the statistical significance of *δ*(*t*) and the sign (improved vs. impaired growth) differs between batches (Fig 4B, left). A similar batch variability was observed under high OS, but due to the stronger effect of the stress perturbation, estimates of *δ* (*t*) are less affected by batch and replicate variation (Fig 4B, right). Similarly, the batch variability observed in the raw *P. aeruginosa* growth data (Fig. 1B) results in significantly different posterior estimates of the interaction effect (*αβ*)_*p,m*_(*t*) across batches (Fig. 4C). Differences observed include the timing and length of negative growth impact (benzoate and citric acid), and completely opposite effects with either strong or no interaction (malic acid). In addition, the posterior variance of each function, which indicates the level of uncertainty remaining, is low for each batch modeled separately. This indicates high confidence in the estimated function despite observed differences across batches. These analyses suggest that use of a single experimental batch leads to overconfidence in explaining the true underlying growth behavior.

### 2.3 Hierarchical models correct for batch effects in growth data

To demonstrate the use of *phenom* to combat the impact of batch effects on growth curve analysis, we combined data across all batches and performed the analysis using each of the M_null_, M_batch_, and M_full_ models (Fig. 5). Estimates of *µ*(*t*) between each model were largely similar, likely due to the abundance of data present to estimate this variable (Fig. S6). Instead, we focus on the estimates of *δ*(*t*) for low and high OS response of *H. salinarum* (Fig.5A) *and the interaction (αβ*)_*p,m*_ between pH and OA concentration effects on *P. aeruginosa* growth (Fig. 5C).

**Figure 5:**
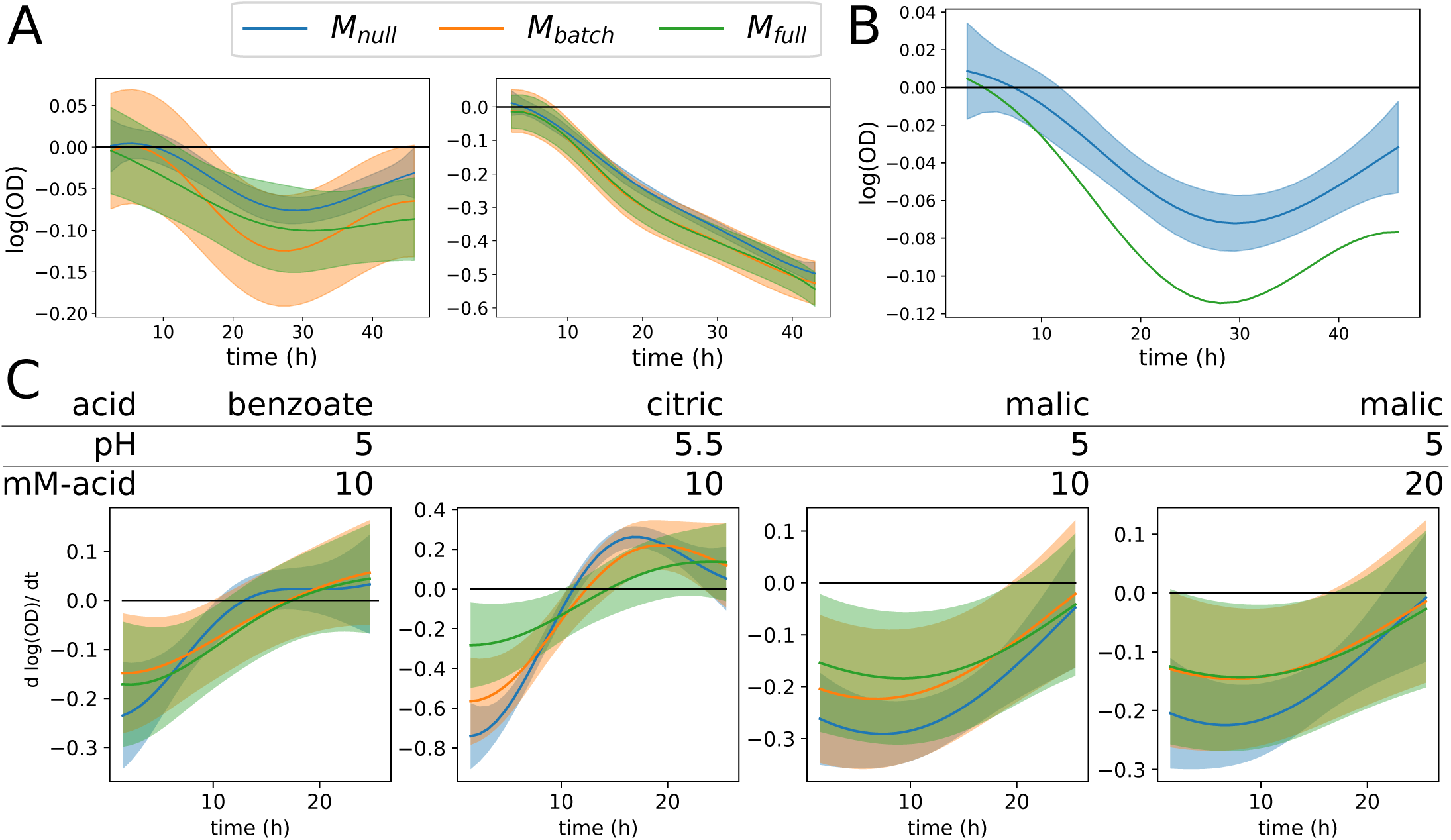
Hierarchical models of growth control for batch effects. Posterior intervals of functions estimated by models of increasing hierarchical complexity: M_null_ (blue), M_batch_ (orange), and M_full_ (green). Solid line indicates posterior mean and shaded regions indicate 95% credible regions. (A) Posterior interval of *δ* (*x*) for low (left) and high (right) OS response by *H. salinarum*. (B) Posterior interval of *δ* (*x*) under M_null_ (blue shaded region) compared to the posterior mean of M_full_(green line). (C) Posterior interval of interaction function (*αβ*)_*p,m*_(*t*) for *P. aeruginosa* growth in indicated pH and organic acid concentration.

Growth impairment in the presence of low OS relative to standard conditions (i.e. *δ* (*t*)) is estimated to be significant during the time points of ∼ 10 − 40 hours under M_null_. In contrast, only time points ∼ 20 − 40 are significantly non-zero under M_batch_ (Fig. 5A, left). Although M_full_ and M_null_ exhibit similar regions of time where effects are significant, uncertainty is higher (confidence bands wider) when batch and replicate effects are taken into account (M_full_). Given the stronger stress effect in the high OS condition (Fig. 5A, right), estimates of *δ* (*t*) were significantly non-zero under all three models, with only minor differences between the three model estimates. Importantly, we note that the posterior interval of *δ* (*t*) under M_null_ for low OS does not include the best approximation of the true function (the posterior mean of *δ* (*t*) under M_full_) for greater than 80% of the time course (Fig. 5B). Taken together, these results suggest that certain time points where *δ* (*t*) is concluded to be non-zero under M_null_ may be inaccurate, especially for stress conditions with modest effects on growth phenotype.

The impact of modeling hierarchical variation on estimating interaction effects in *P. aeruginosa* growth was condition dependent (Fig. 5C). Across conditions, however, a decrease in posterior certainty on the true shape of the underlying function was again observed under M_batch_ and M_full_. For example, the interaction between benzoate and pH became less pronounced under M_full_. Similarly, the models of (*αβ*)_*p,m*_(*t*) under citric and malic acid showed shrinkage toward zero under M_batch_ and M_full_. Such shrinkage is a common observance in hierarchical modeling^53^. Taken together, these results for *P. aeruginosa* extend those previously published^23^, which only included analysis using the M_null_ model.

For both *H. salinarum* response to OS and *P. aeruginosa* growth under pH and OA exposure, an increase in posterior variance was observed under M_batch_ and M_full_ compared to M_null_ (Fig S7). However, posterior variance of *δ* (*t*) in the *H. salinarum* OS response was higher under M_batch_ compared to M_full_. In this case, controlling for replicate effects appears to increase the signal needed to identify *δ* (*t*). In contrast, these variances are equal in the *P. aeruginosa* data, indicating that the relative improvement in variance afforded by modeling batch vs. replicate effects may be dataset dependent.

### 2.4 Variance components demonstrate the importance of controlling for batch effects

Variance components, which correspond to the estimated variance of each effect in the model, can be used to compare the impact each group has on the process of interest^24^. To better understand sources of variability in growth curve studies, we used *phenom* to estimate the variance components for each dataset above. In our hierarchical non-parametric setup, these variance components are the variance hyperparameters (e.g. *σ*^2^) of the Gaussian process kernels for each fixed and random effect group. These parameters control the magnitude of function fluctuations modeled by the GP distribution. Larger variance implies higher effect sizes and therefore a larger impact on the observations.

We show the value of variance components by considering the effects identified by M_full_ for *H. salinarum* under low OS (Fig. 6). The variance of the data is partitioned between the mean growth (*µ*(*t*)), the OS (*δ*(*t*)), batch effects (batch curves of *µ*(*t*) and *δ*(*t*)), biological noise (e.g. replicate variability) and instrument noise 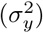. This analysis confirms that batch effects, compared to the other sources of experimental variability in the dataset (replicate noise and measurement error), are between 2 to 10 times more impactful on the phenotype measurements. Additionally, variance components enable comparisons between the experimental and treatment factors in the data. Of particular note is that the variance of the treatment of interest, *δ* (*t*), and the batch effects are similar in magnitude, at least in the case of a low-magnitude stress such as 0.083 PQ for *H. salinarum*. This suggests that proper modeling of this treatment requires both sufficient batch replication and accurate modeling of batch effects in those data. Future studies of similar phenotypes can be guided by these estimates in experimental design, choosing an appropriate batch replication for the degree of noise expected^57^. However, the extent of replication required may depend upon the dataset (factorial design, treatment severity, etc). Taken together, variance components provide an aggregated view of the contribution by various factors and guide future experimentation.

**Figure 6:**
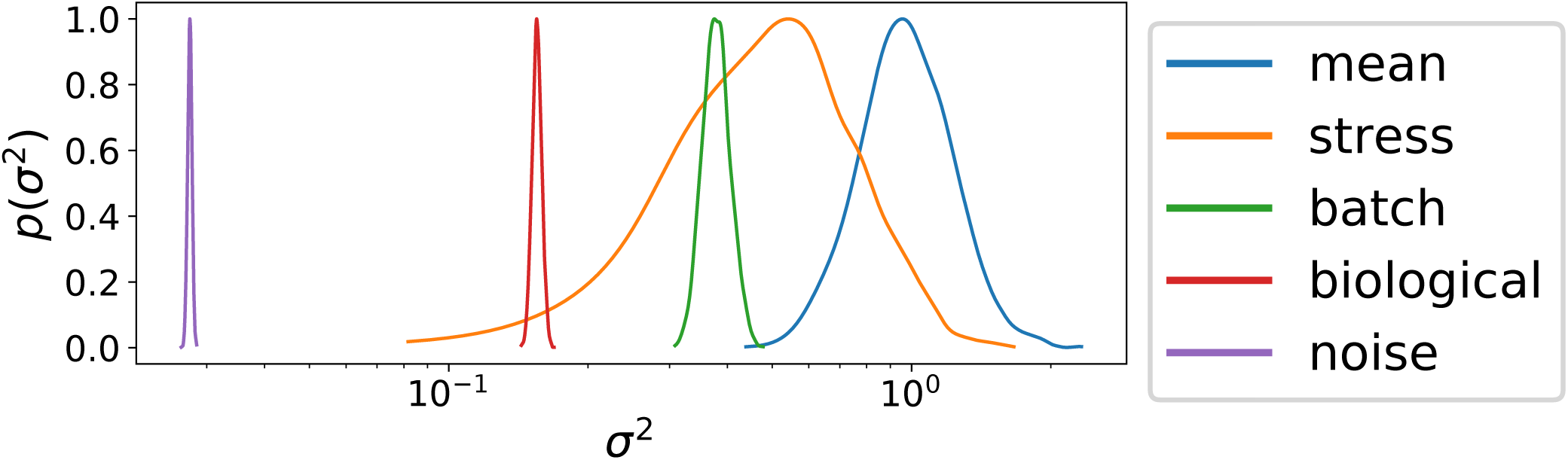
Posterior variance components in the *phenom* hierarchical phenotype model. Posterior intervals are shown for the kernel variance hyperparameter for different groups of effects from *phenom* estimated on *H. salinarum* growth under low OS. Groups correspond to *µ*(*t*) (mean), *δ* (*t*) (stress), batch effects (batch), replicate noise (biological), and measurement error (noise).

## 3 Discussion

We have provided a framework to test and control for random effects in microbial growth data using the hierarchical non-parametric Bayesian model, *phenom* (Fig. 3). Analysis with *phenom* indicates that random effects (both batch and replicate) appear in the two microbial population growth datasets studied here, and constitute significant portions of the variability (Fig. 1). Failure to correct for these effects confounds the interpretation of growth phenotypes for factors of interest in a large scale phenotyping analysis (Fig. 4). *phenom* controls for these random effects and provides accurate estimates of the growth behavior of interest (Fig. 5). Additionally, *phenom* can be used to estimate variance components, providing information about the relative impact of various sources of noise in the data (Fig. 6). Controlling for batch effects in these datasets was therefore key to making accurate biological conclusions.

Related fields of functional genomics, such as transcriptomics, have seen considerable interest in controlling for different experimental sources of variation, broadly labeled as batch effects^28,57–62^. These studies have shown that differences between batches first need to be corrected to avoid erroneous conclusions^63^. Here we have shown that, like in transcriptomics data, controlling for sources of variation in phenomics data - particularly due to batch - are an important step in making accurate biological conclusions regarding population growth.

*phenom* establishes a complete and general method of controlling batch effects in microbial growth phenotypes, overcoming significant weaknesses of previously developed techniques. In reference [19] we identified and corrected for batch effects in a single transcription factor mutant’s stress response, but this model did not provide an explicit deconstruction of batch effects between different factors (e.g. strain and stress) and could therefore not determine which factors were most strongly impacted by batch effects. Moreover, this approach utilized a standard GP regression framework, but the standard framework has well-established limitations on dataset size, limiting its applicability to the large datasets we consider here. In reference [22] we described a functional ANOVA model for microbial growth phenotypes, which corresponds to the M_null_ model in the *phenom* case. Again, a global batch effects term was included but individual batch effects were not modeled, and the computational approach utilized (Gibbs sampling) was prohibitively slow for the complete *phenom* model.

Although we focus here on replicate and batch variation, the *phenom* model is easily extended to incorporate alternative or additional random and fixed effects appropriate for settings with other sources of variation. For example, depending on the experimental design, *phenom* could control for variation among labs, experimental material, culture history, or genetic background^25,64–70^. *phenom* flexibly incorporates additional sources of variation and/or interaction between design variables, as demonstrated with the two different designs analyzed for *H. salinarum* and *P. aeruginosa* here. This flexibility allows *phenom* to be applied to control for many sources of technical variation within microbial population growth data, thereby improving the analysis and resulting conclusions regarding quantitative microbial phenotypes.

## 4 Materials and Methods

### 4.1 Experimental Growth Data

*H. salinarum* growth was performed as described previously^22^. Briefly, starter cultures of *H. salinarum* NRC-1 Δ*ura*3 control strain^71^ were grown at 42°C with shaking at 225 r.p.m. to an optical density at 600 nm (OD_600_) ∼ 1.8 − 2.0 in 3 mL of Complete Medium (CM; 250 NaCl, 20 g/l MgSO4•7H2O, 3 g/l sodium citrate, 2 g/l KCl, 10 g/l peptone) supplemented with uracil (50 *µ*g/ml). Cultures were then diluted to OD_600_∼ 0.05 in a high throughput microplate reader (Bioscreen C, Growth Curves USA, Piscataway, NJ), and growth was monitored automatically by OD_600_ every 30 minutes for 48 hours at 42°C. High and low levels of OS were induced by adding 0.333 mM and 0.083 mM of paraquat to the media, respectively, at culture inoculation.

For *P. aeruginosa*, laboratory strain PAO1 (ATCC 15692) was grown as described in reference [23]. Briefly, cultures were grown in M9 minimal media supplemented with 0.4% (w/v) glucose and 0.2% (w/v) casamino acids and buffered with 100 mM each of MES and MOPS buffers. Population growth was measured with a CLARIOstar automated microplate reader (BMG Labtech) at 37°C with 300 rpm continuous shaking. The OD_600_ was recorded automatically every 15 minutes for a total of 24 hours. A full factorial design of pH and OA concentration was performed for benzoate, citric acid, and malic acid. An experimental batch corresponded to two repetitions of the experiment on separate days with a minimum of three biological replicates of each condition on each day. Two batches for each OA were performed.

All data generated or analysed during this study are included in this published article (see supplementary information files).

### 4.2 Parametric growth curve estimation

For comparison with our non-parametric methods, parametric growth curve models were estimated using the grofit package in R with default parameters^72^. The logistic model was used to fit each curve. Kernel density estimates of parameter distributions were calculated with the scipy package with default kernel bandwidth parameters^73^.

### *4.3 phenom*: a hierarchical Gaussian process model of microbial growth

#### 4.3.1 Gaussian Processes

A Gaussian process (GP) defines a non-parametric distribution over functions *f* (*t*), defined by the property that any finite set of observations of *f* follow a multivariate normal distribution^54^. A GP is fully defined by a mean function *m*(*t*) and a covariance function *κ*(*t, t*′):

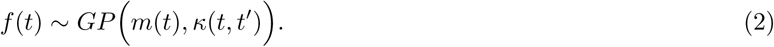

GPs are commonly used for non-parametric curve fitting^54^ where *m*(*t*) is typically set to 0, which we do here. Similarly, we use a common choice for covariance function defined by a radial basis function (RBF) kernel:

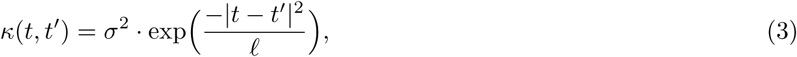

where *σ*^2^ is the variance and *ℓ* is the length-scale. The parameter *σ*^2^ controls the overall magnitude of fluctuation in the population of functions described in the GP distribution, while *ℓ* controls the expected smoothness, with larger *ℓ* making smoother, slower varying functions more likely. In the process of non-parametric modeling of growth curves, these parameters are adaptively estimated from the dataset.

#### 4.3.2 Fixed effects

We first define the fixed effects models used in this study; these will be augmented with random effects in the next section. We consider fixed effects models of increasing complexity: a mean growth phenotype, a single treatment phenotype, and a combinatorial phenotype with interactions between treatments. All of these models fall under the functional analysis of variance (ANOVA) framework^22,74^. To estimate a mean growth profile, as in the case of measuring a single condition, a mean function *µ*(*t*) is estimated from the data by modeling each replicate *y*_*r*_(*t*) for 1 ≤ *r* ≤ *R* as consisting of an unknown mean function observed with additive noise:

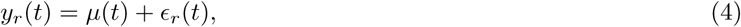

where *µ*(*t*) ∼ *GP* (0, *κ*_*µ*_(*t, t*′)) provides a prior distribution over *µ*, and *κ*_*µ*_ is an RBF kernel with hyperparameters 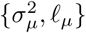. Here 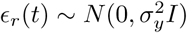 is Gaussian white noise.

When estimating the effect of a perturbation on growth, as in the case of OS, we add a second function *δ* (*t*) that represents the effect of the stress being considered. The model then becomes

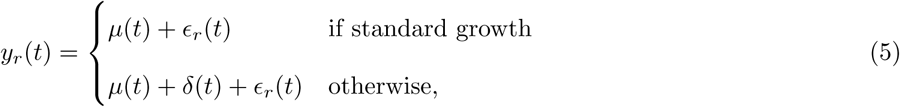

where *δ* (*t*) ∼ *GP* (0, *κ*_*δ*_ (*t, t*′)) also follows a GP prior independently of *µ*, and *κ*_*δ*_ has hyperparameters 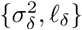.

When incorporating possible interaction effects such as those between pH and organic acids in the *P. aeruginosa* dataset, the model becomes

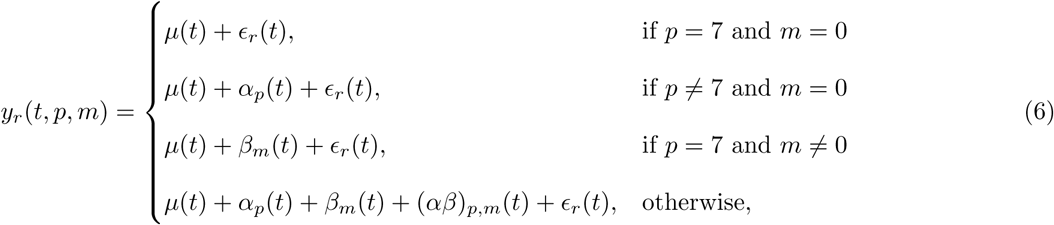

for pH *p* and molar acid concentration *m*, with *α*_*p*_(*t*) representing the main effect of pH, *β*_*m*_(*t*) the main effect of acid concentration, and (*αβ*)_*p,m*_(*t*) the interaction between them. Each effect is drawn from a treatment specific GP prior:

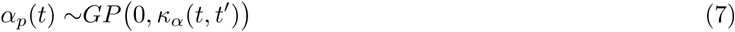

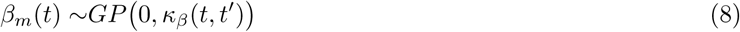

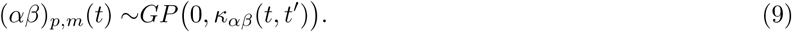

Again, each covariance function is specified by a RBF kernel with corresponding variance and lengthscale hyperparameters that adapt to the observed data. All models in this section correspond to M_null_ for their respective analyses, as they do not include any random effects.

#### 4.3.3 Random effects

The first random effects added to the model were those used to account for batch effects, in the model M_batch_. Under this model, each fixed functional effect becomes the mean of a GP describing the population of possible batch-specific mean curves. For example, under the model of mean growth behavior (Eq. 4), replicate *r* from batch *i* is modeled as

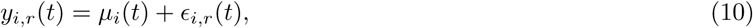

where *µ*_*i*_ is the batch mean drawn from *µ*_*i*_(*t*) ∼ *GP* (*µ*(*t*), *κ*_*µ*,batch_(*t, t*′)) with kernel *κ*_*µ*,batch_ and 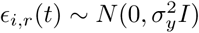. Other M_null_ models are converted to M_batch_similarly, with each fixed effect becoming a mean of a GP prior for each batch effect. M_full_ develops the hierarchy one step deeper by adding replicate effects to M_batch_. Specifically, the error model *ϵ*_*i,r*_ is now described by a GP: *ϵ*_*i,r*_ ∼ *GP* (0, *κ*_*y*_(*t, t*′)) with corresponding hyperparameters, accounting for replicate-specific variability rather than simply white noise.

#### 4.3.4 Inference

As noted above, each group GP prior is specified by its own RBF kernel with corresponding variance and length-scale parameters 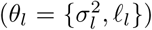. For each group, 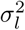 is assigned a Gamma(*α, β*) prior and *ℓ* _*l*_ a conjugate inverse-Gamma prior, with user-defined hyperparameters. Noise variance 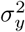 was also assigned a gamma prior. Bayesian inference was then performed, with the posterior distribution obtained by sampling using Markov chain Monte Carlo (MCMC) implemented with the Stan library, which uses a Hamilitonian Monte-Carlo procedure with No-U-turn sampling^55^. Multiple chains were run to diagnose convergence, with all parameter posterior means confirmed to have converged within 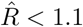 as recommended^75^.

### 4.4 Data and Code Availability

All code for this study is available at https://github.com/ptonner/phenom. Raw growth data are available for *H. salinarum* in reference [22] and at https://github.com/ptonner/hsalinarum_tf_phenotype. Raw growth data are available for *P. aeruginosa* in reference [23] and at https://github.com/amyschmid/pseudomonas-organic-acids.

## 5 Acknowledgements

## 5.1 Funding

This study was funded by National Science foundation grants DMS-1407622 to SCS; NSF-MCB-1615685, −1651117, and −1642283 to AKS; NSF Graduate Research Fellowship to PDT. Research on this project in PAL’s laboratory was supported by grant number BB/K019171/1 from the UK Biotechnology and Biological Sciences Research Council.

## 5.2 Author Contributions

PT, AS, PL, and SS conceived of the study. AS, SS, PL directed and provided oversight, training, and funding for the study. PT performed analysis and generated software. CD and FB performed experiments. PT, AS, and SS wrote the manuscript. All authors contributed to the final draft.

## 6 Supplementary Material

**Figure S1:**
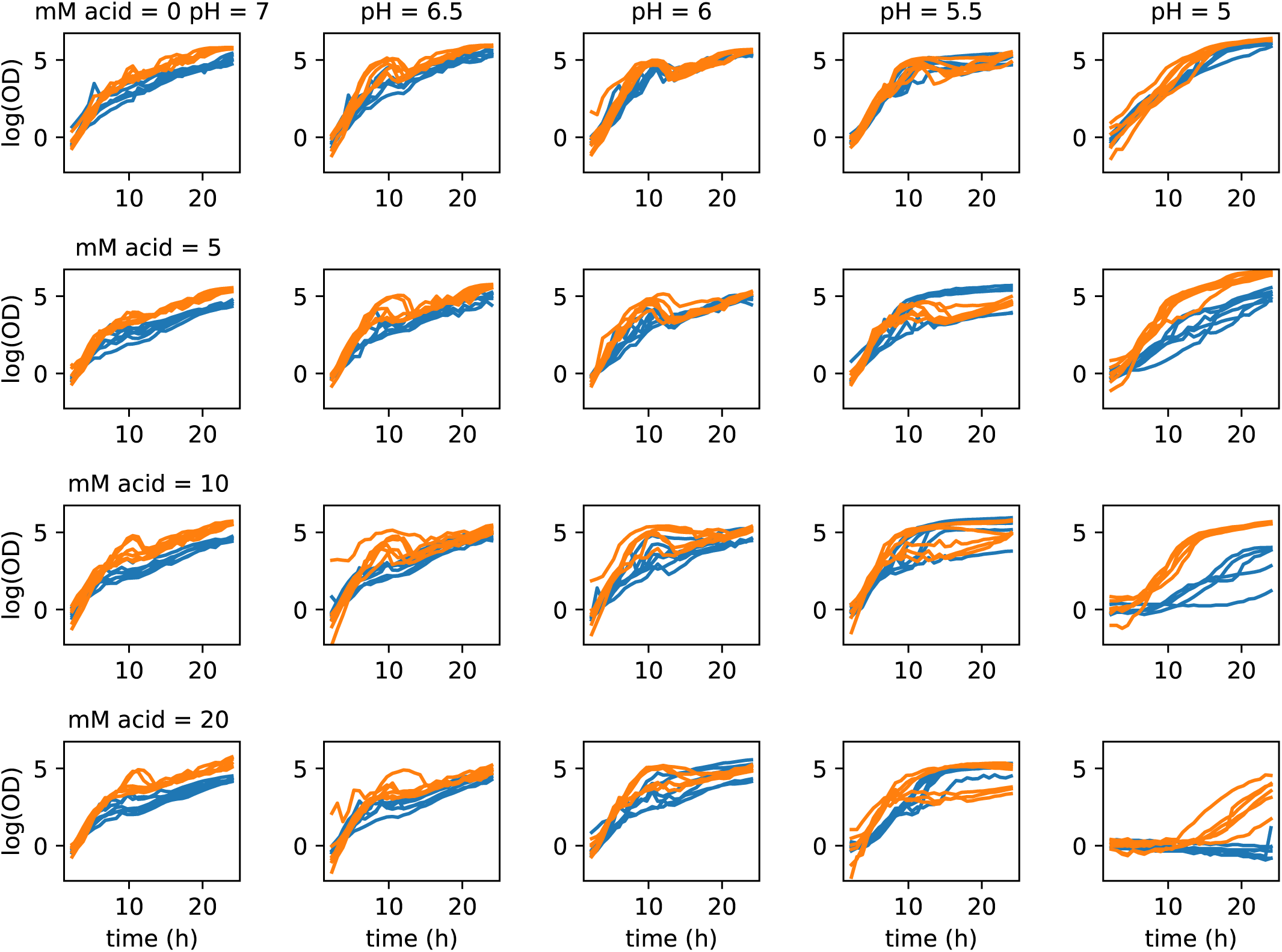
*P. aeruginosa* growth under benzoate and pH gradient. Growth of *P. aeruginosa* strain PA01 under gradient of pH (7 — 5) and benzoate (0 — 20). Colors represent different batches.

**Figure S2:**
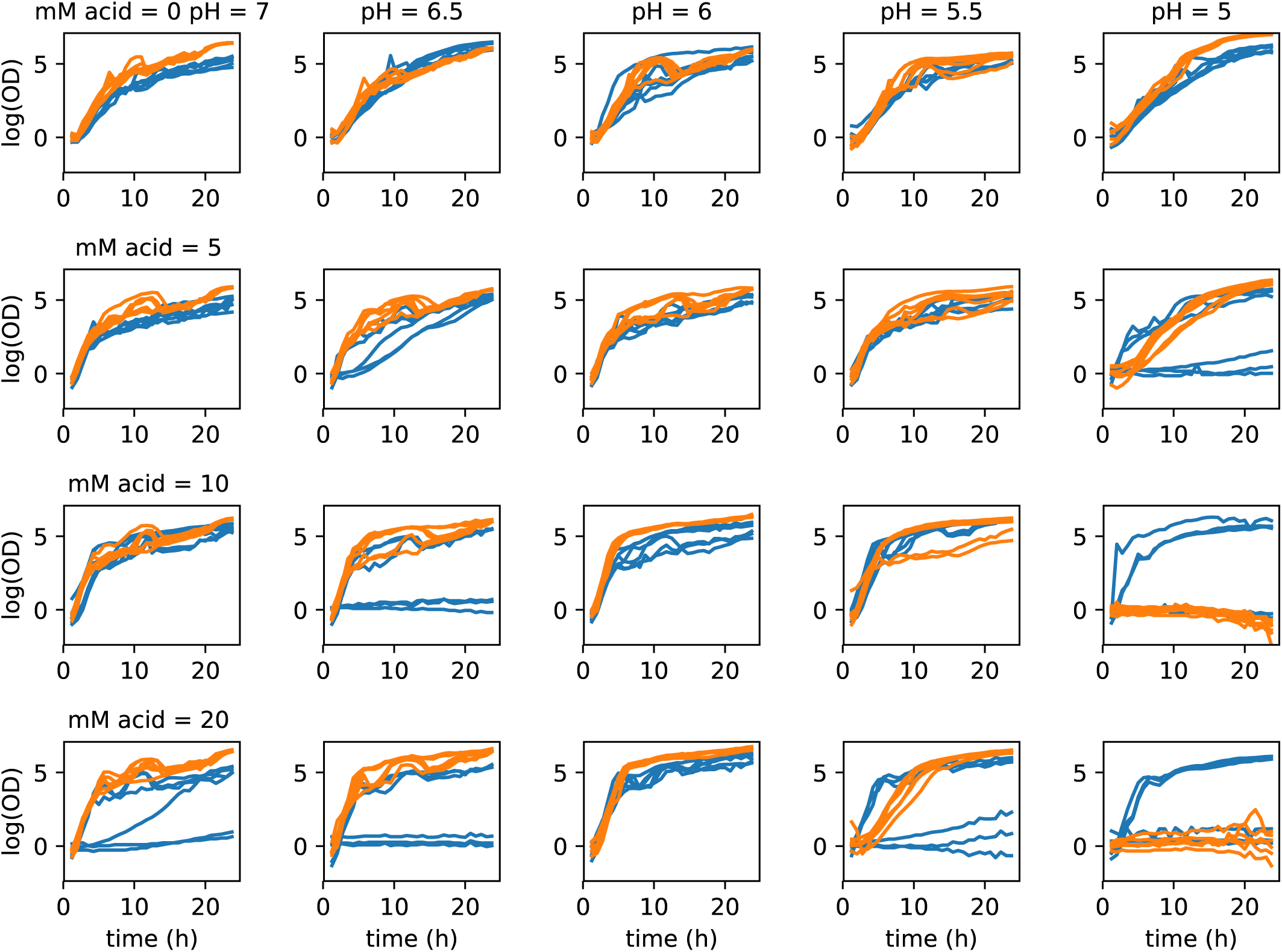
*P. aeruginosa* growth under malic acid and pH gradient. Growth of *P. aeruginosa* strain PA01 under gradient of pH (7 — 5) and malic acid (0 — 20). Colors represent different batches.

**Figure S3:**
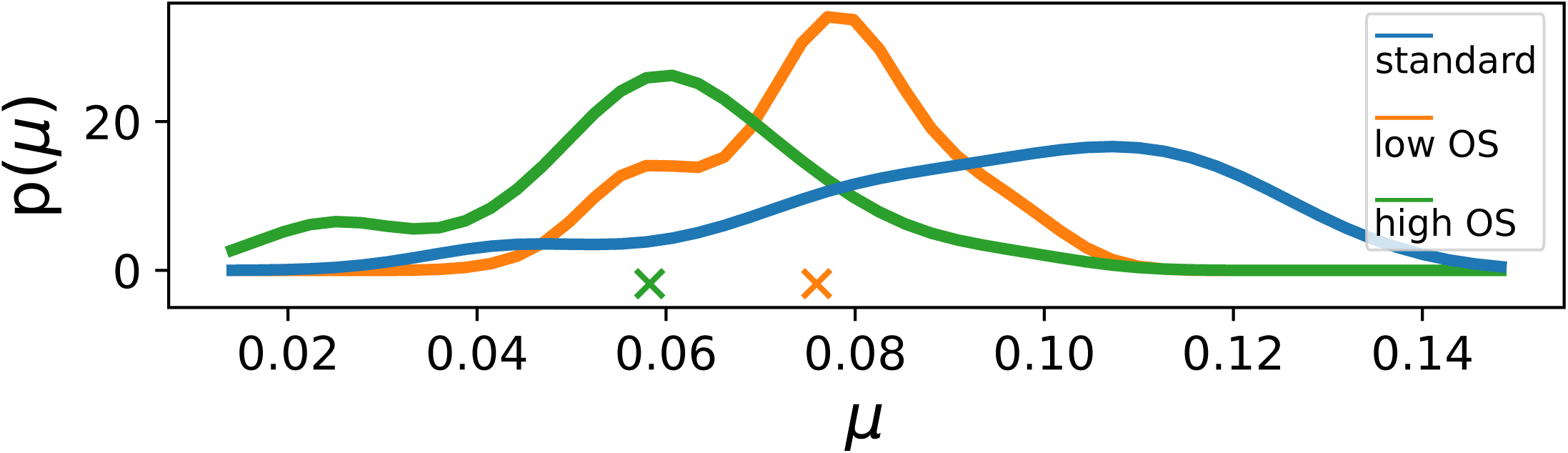
KDE of *µ*_max_ for *H. salinarum* growth across batches. Crosses indicate significant difference between *µ*_max_ standard conditions and each OS level (one-sided t-test, *p <* 0.05)

**Figure S4:**
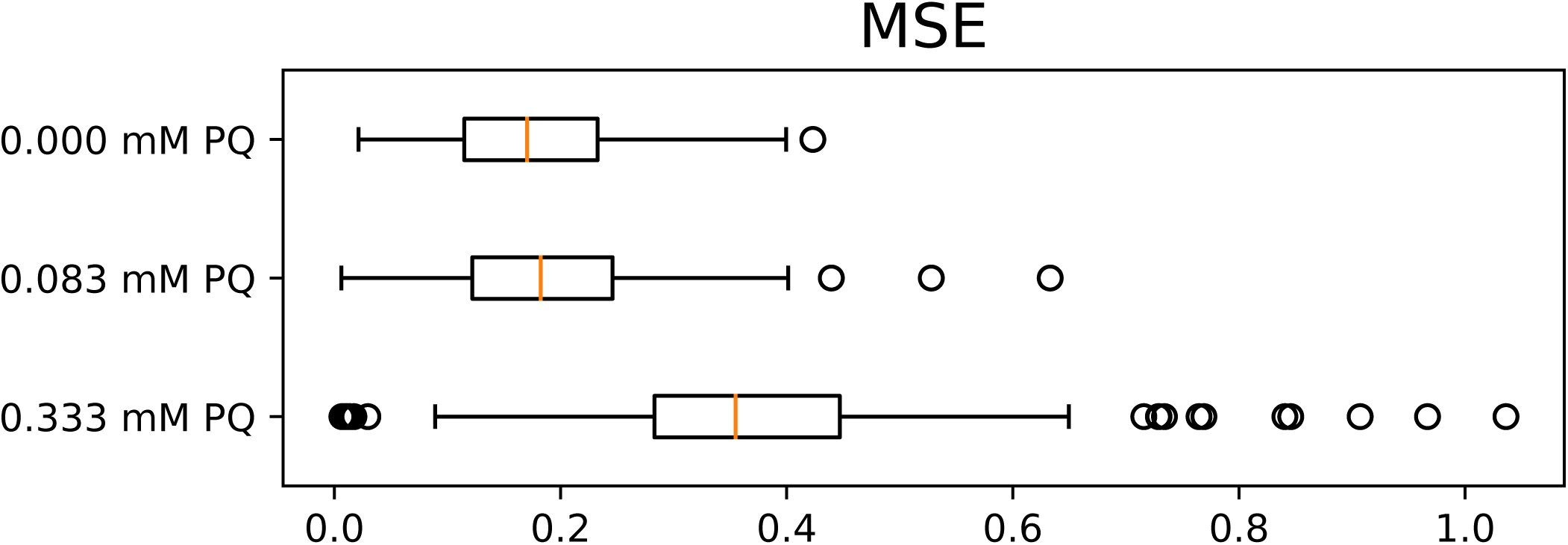
Error in parametric growth models. Distribution of error (MSE) for each condition when fit with a logistic growth curve. The box show shows the inter-quartile range, red line is the median, whiskers show the 1.5 inter-quartile range, and the individual points are outliers.

**Figure S5:**
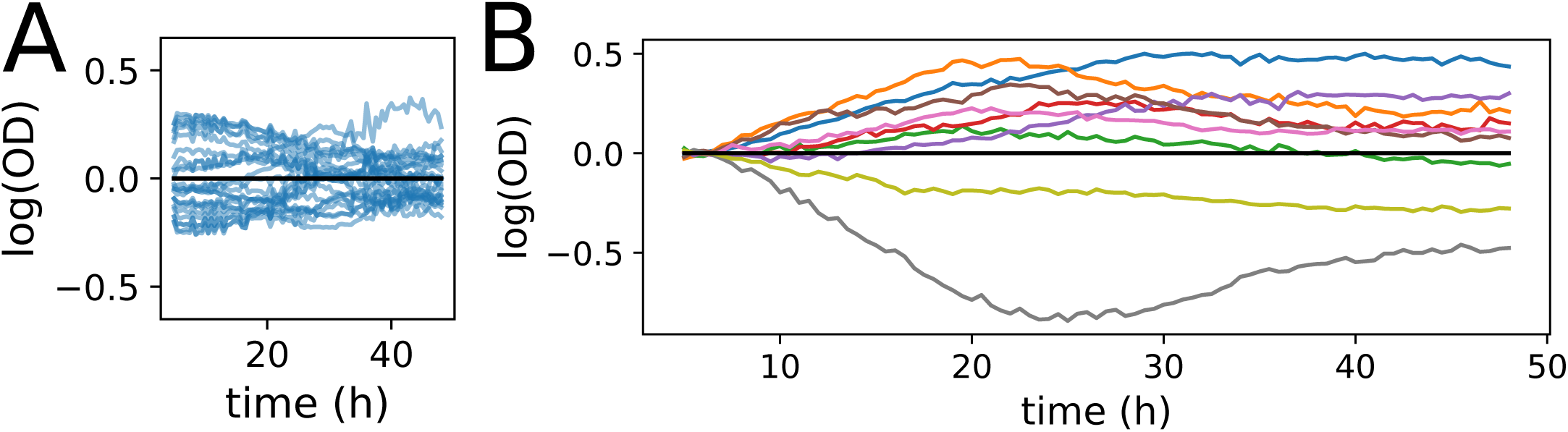
Residual structure of microbial growth data across batches. (A) Individual replicate curve residuals around the mean of the respective batch. Only standard conditions are shown. (B) Residual of the mean behavior for each batch around the global mean (standard condition only).

**Figure S6:**
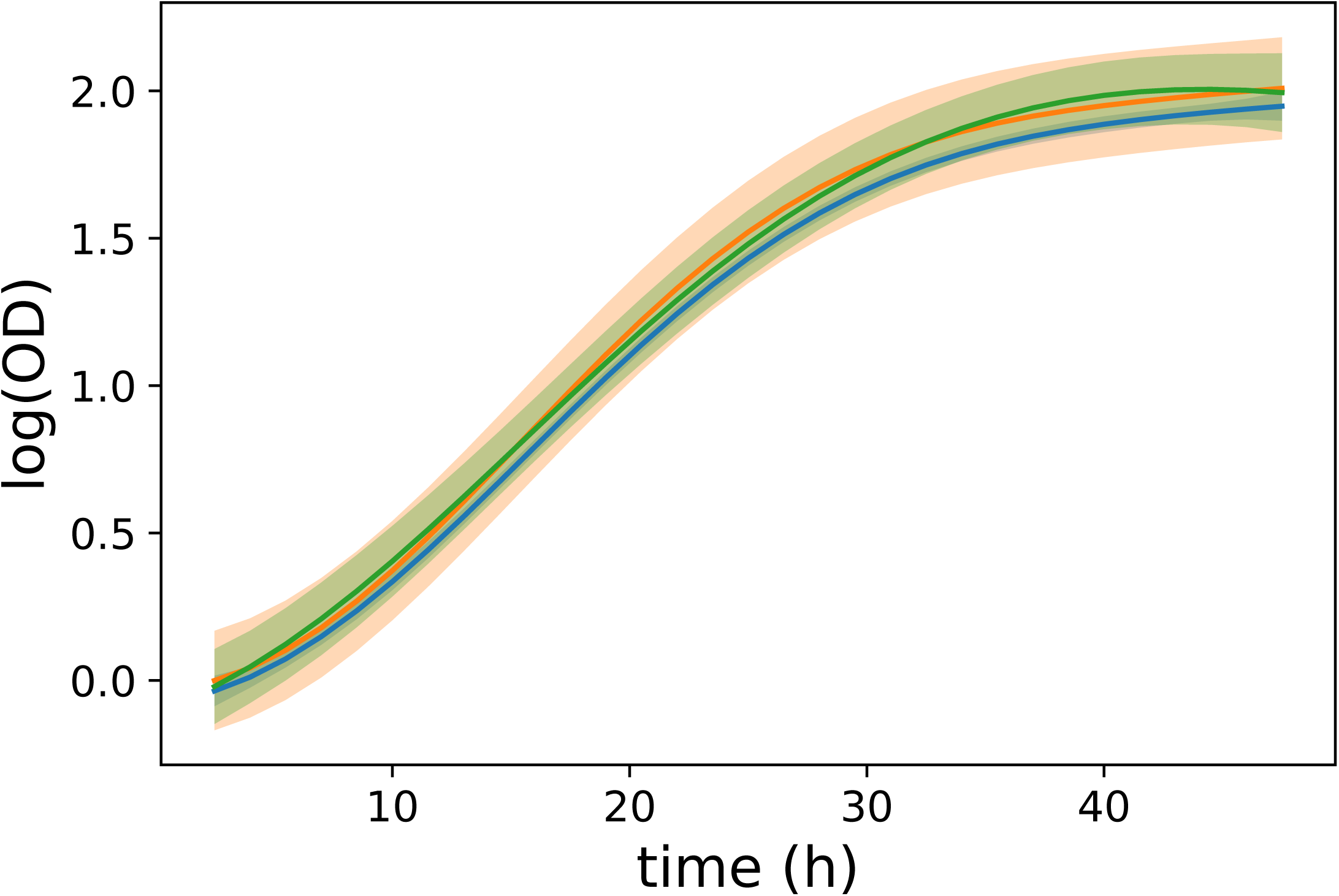
Posterior comparison of *µ*(*t*) for *H. salinarum* growth across batches. Posterior interval of *µ*(*x*) for *H. salinarum* standard growth.

**Figure S7:**
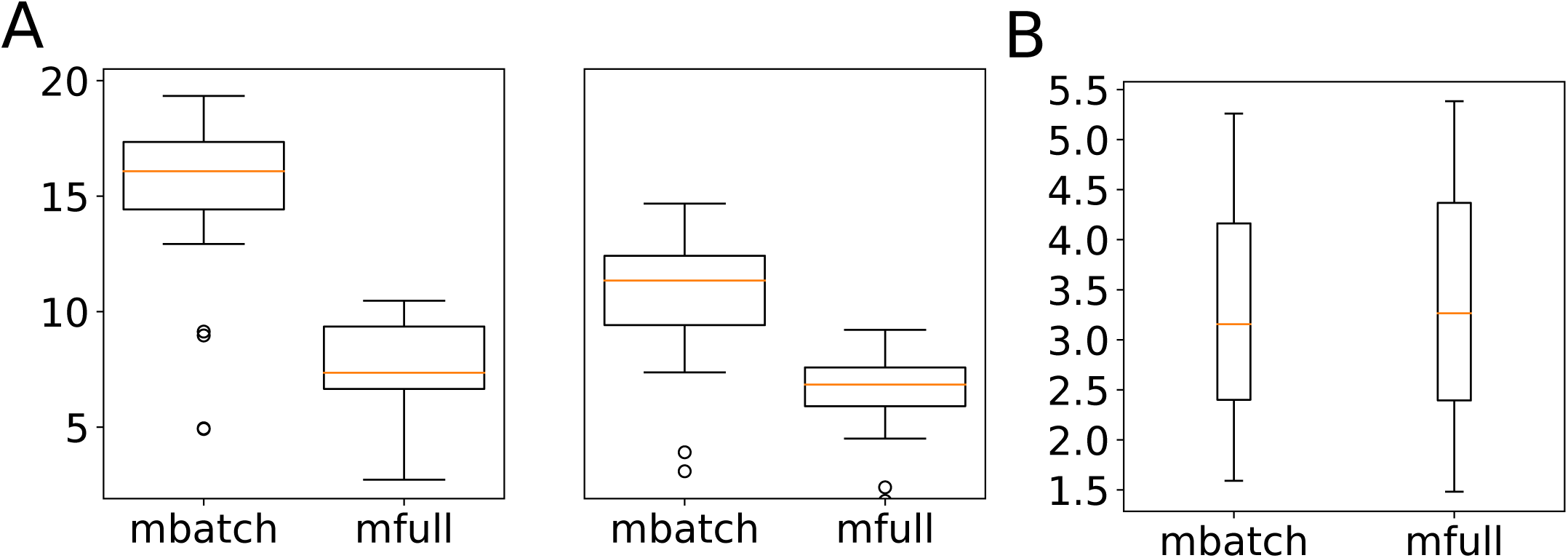
Posterior variance of function estimates under different models. Each plot shows the posterior variance of a function at each time point under each of M_batch_ and M_full_ versus M_null_. (A) *δ* (*x*) estimated for *H. salinarum* growth under low (left) and high (right) OS. (B) (*αβ*)_*p,m*_(*t*) at pH = 5, mM malic acid = 10.

## Notes

https://github.com/ptonner/phenom

